# Phenogenomic resources immortalized in a panel of wild-derived strains of five species of house mice

**DOI:** 10.1101/2023.11.05.565684

**Authors:** Jaroslav Piálek, Ľudovít Ďureje, Zuzana Hiadlovská, Jakub Kreisinger, Tatiana Aghová, Anna Bryjová, Dagmar Čížková, Joëlle Goüy de Bellocq, Helena Hejlová, Kateřina Janotová, Iva Martincová, Annie Orth, Jana Piálková, Iva Pospíšilová, Ludmila Rousková, Barbora Vošlajerová Bímová, François Bonhomme, Jiří Forejt, Miloš Macholán, Pavla Klusáčková

## Abstract

The house mouse, *Mus musculus*, is a widely used animal model in biomedical research, with classical laboratory strains (CLS) being the most frequently employed. However, the limited genetic variability in CLS hinders their applicability in evolutionary studies. Wild-derived strains (WDS), on the other hand, provide a suitable resource for such investigations. This study quantifies genetic and phenotypic data of 101 WDS representing 5 species, 3 subspecies, and 8 natural Y consomic strains and compares them with CLS. Genetic variability was estimated using whole mtDNA sequences, the *Prdm9* gene, and copy number variation at two sex chromosome-linked genes. WDS exhibit a large natural variation with up to 2173 polymorphic sites in mitogenomes, whereas CLS display 92 sites. Moreover, while CLS have two *Prdm9* alleles, WDS harbour 46 different alleles. Although CLS resemble *M. m. domesticus* and *M. m. musculus* WDS, they differ from them in 8 and 11 out of 15 phenotypic traits, respectively. The results suggest that WDS can be a useful tool in evolutionary studies and have great potential for medical applications.

## Introduction

Progress in many areas of the life sciences would be inconceivable without appropriate biological models. For example, the *Drosophila* fruit fly was used to establish the fundamentals of genetics ^1,2^, while yeast was used to discover DNA recombination mechanisms and gene-protein functions ^3^. The laboratory mouse is the most prominent vertebrate model ^4^. Mice are easy to keep, breed, and handle, they are also highly tolerant to inbreeding, and their normative biology is well understood ^5^. A whole-genome sequence was published almost simultaneously with that of humans ^6^, and large databases of well-annotated sequences, SNPs, and other markers with known positions in the genome are now available, making the laboratory mouse an ideal organism for tracing genotype-phenotype associations and inferring gene function e.g., ^7^. The ability to replenish breeding stocks from frozen embryos avoids the negative effects of mutation load and thus increases the reproducibility of experimental preclinical studies. Due to a relatively short generation time and the ability to reproduce throughout the year, it is possible to generate multiple laboratory stocks within a reasonable time. In fact, dozens of inbred ‘classical laboratory strains’ (CLS) have been established since the early 20^th^ century. There are more than 200 CLS listed in ^8^, indicating the availability of significant genetic diversity that can be harnessed when selecting the most suitable mouse model for a specific study.

However, despite the indispensability of CLS for biomedical and life science studies, their applicability appears to be reaching its limits. First, their genome is an artificial mixture of three house mouse subspecies: *Mus musculus domesticus*, *M. m. musculus*, and *M. m. castaneus* ^9,10^. Second, despite the high number of existing CLS, they display limited genetic variation. For example, they all essentially represent a single female as demonstrated by Ferris et al. ^11^ and later confirmed by mitochondrial whole-genome sequences ^12–14^ or hundreds of thousand SNP in over 150 CLS ^9^. This finding poses a significant limitation when one wishes to apply CLS-based genotype-phenotype associations to human populations that exhibit high genetic variability and, subsequently, an increased level of (epi)genetic interactions. Naturally, the highest variation in house mice occurs within wild populations. While geographically wide-scale whole genome sequencing of wild mice has not progressed as extensively as human studies (as exemplified by the 1000 Genomes Project or the Pangenome Reference project ^15,16^), recent publications have summarized diversity across more than 310 shotgun-read mouse genomes ^17–26^. This diversity is further expanded by introgressive hybridization observed in their contact zones, such as between *M. m. castaneus* and *M. m. musculus* ^22^ or *M. m. domesticus* and *M. m. musculus* subspecies ^27,28^. All these studies reveal that each mouse represents a unique genotype. In contrast to inbred laboratory stocks, the high natural variation in wild mice conflicts with strict requirements for experimental reproducibility. Moreover, the inevitable presence of rare alleles in wild populations hampers genome-wide association studies (GWAS) ^29,30^.

An excellent compromise between the two contradicting requirements, i.e. availability of suitable variation and reproducibility, appears to be inbred wild-derived strains (WDS) ^26,31–33^. Contrary to CLS, their geographic origin and pedigree are precisely known. In addition, inbred WDS are thought to represent immortal resources suitable for GWAS as the rare alleles are fixed for different variants in individual strains. Although WDS cannot fully capture natural variation, this deficiency can be reduced by increasing the number of WDS used. Thus, by exploiting their phenotypic variation and genomic complexity, WDS provide an ultimate model system for understanding the genetic control of quantitative traits of biomedical and evolutionary interest. To address these questions, we estimated genetic and phenotype variability in more than 100 WDS representing five species of the genus *Mus* and compare it with CLS. We analyzed whole-genome mitochondrial (mtDNA) sequences, partial sequences of the *Prdm9* gene known to cause reproductive isolation between the *M. m. domesticus* C57BL/6J (CLS) and *M. m. musculus* PWD (WDS) strains ^34,35^, copy number variation in two sex chromosome-linked genes, and a set of phenotypic characters (external morphological traits, reproductive organs in both sexes, and reproductive performance). We show that WDS differ significantly from CLS in genetic and phenotypic variation. The combination of variability preserved in WDS and classic lab mice can be the basis for creating a robust house mouse model not only for evolutionary but also for biomedical research.

### Data Records

#### Mouse repository

The repository has been established by merging three WDS resources based in Montpellier, France ^36,37^, Prague ^38^, and Studenec, Czech Republic ^39,40^. In total, it consists of 101 strains (94 WDS and 7 CLS), all currently kept (76 alive) or were kept in the past (25 became extinct due to reproductive failure or were removed from the breeding scheme). The WDS were classified into groups representing five species, one synanthropic species (*M. musculus*) with three subspecies (*M. m. musculus*, *M. m. domesticus*, *M. m. castaneus*), and four non-synanthropic species (*M. caroli*, *M. macedonicus*, *M. spicilegus*, *M. spretus*). In the following text, we will use the subspecies and species names of mouse taxa (e.g. *musculus* for *M. m. musculus* or *caroli* for *M. caroli*). For comparative purposes addressed in this paper, we define an eighth group comprising all CLS with mixed but predominantly *domesticus* genomes. The spectrum of WDS captures long-term mouse evolution, estimated to be over 5 million years ago since the split of *caroli* from the remaining species under study ^41^. Detailed information on individual strains, including their place of geographic origin, chromosome number, mtDNA and Y (sub)specific ancestry, the presence of the meiotic driver, *t*-haplotype, generation of inbreeding, and reproductive ability over the last four years, is provided in Figure 1 and Supplementary Table S1. Further details can be found at https://housemice.cz/en/strains.

**Figure 1.**
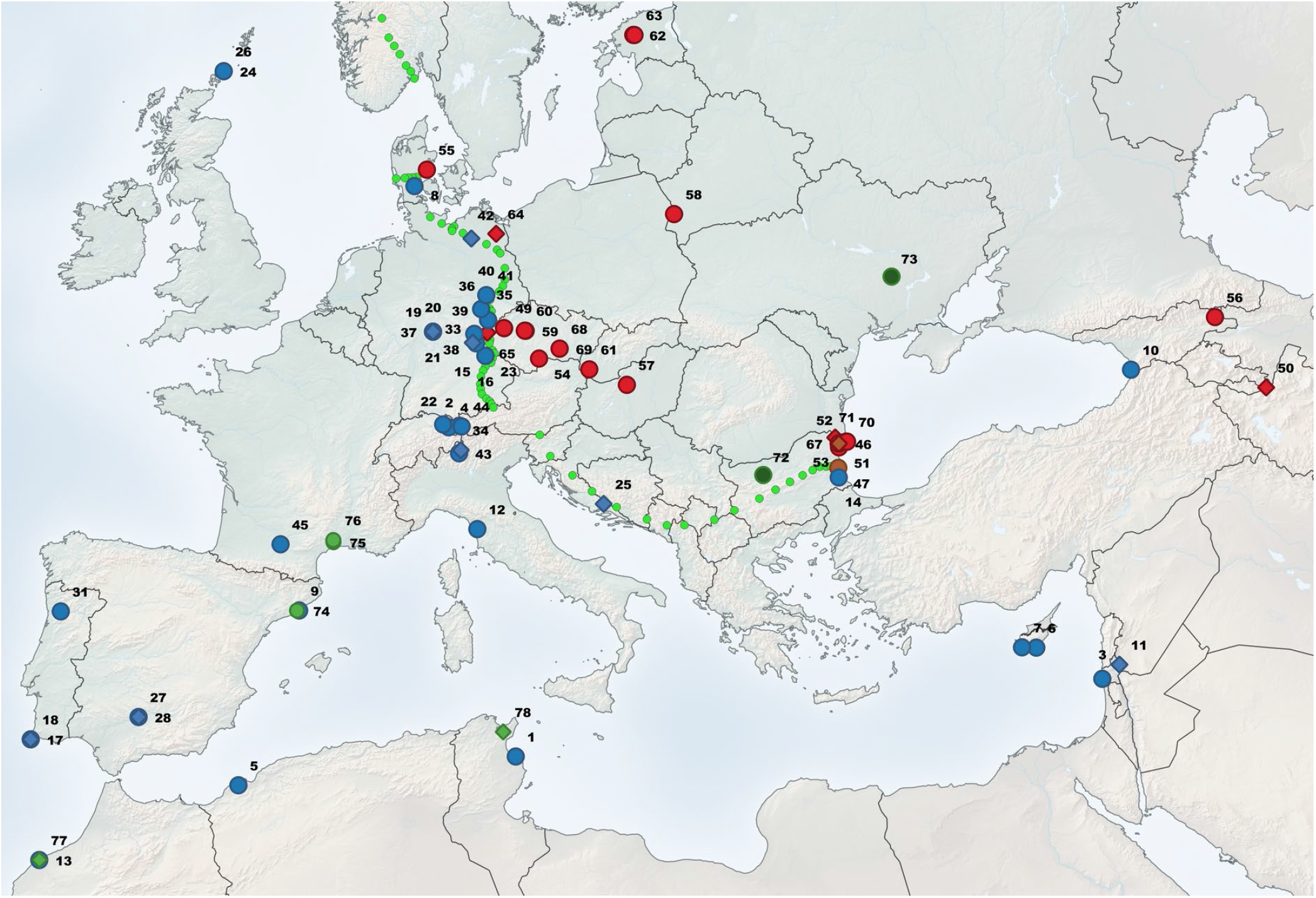
Geographic origin of WDS kept in Studenec; red: *musculus*, blue: *domesticus*, green: *spretus*, brown: *macedonicus*, light olive: *spicilegus*; circles: alive WDS, diamonds: extinct WDS. Small green dots indicate the position of the house mouse hybrid zone adapted from ^28,42^. The numbers labelling localities correspond with those in Supplementary Table S1. WDS out of the depicted area are KTK (*caroli* from Thailand), CIM (*castaneus* from India), DKN and CKN (*castaneus* from Kenya) and MGD (*castaneus* from Madagascar), BID, KAK, and TEH (*musculus* from Iran), MPR (*musculus* from Pakistan), and DOT (*domesticus* from Tahiti, French Polynesia).

#### mtDNA variability

Extra-nuclear variation was assessed using 114 newly sequenced mitogenomes and 49 published whole mtDNA sequences (see Material and Methods and Supplementary Table S2). After alignment, the sequences were 16516 bp long, all representing unique haplotypes, with a total 3,907 polymorphic sites.

Prior to estimating genetic variation, we conducted taxonomic assessment and examined phylogenetic relationships between all the WDS. The maximum-likelihood tree revealed eight distinct clades, each representing a (sub)species. The Madagascan strain (MDG) carried a unique haplotype within the *M. musculus* haplogroups (Figure 2A), similar to the mtDNA lineage previously described as *M. m. gentilulus* ^43,44^. All CLS formed a monophyletic group embedded within the *domesticus* WDS group, reflecting the fact that all CLS carry *domesticus* mtDNA (Figure 2B). The *castaneus* clade consists of two deeply divergent clusters, one involving samples from Iran, and the second from Thailand, India, and Kenya (Figure 2C). This divergence pattern can be attributed either to the presence of a 76-bp insert in the former cluster, or paraphyly of the subspecies as suggested earlier ^45–47^, or both.

**Figure 2.**
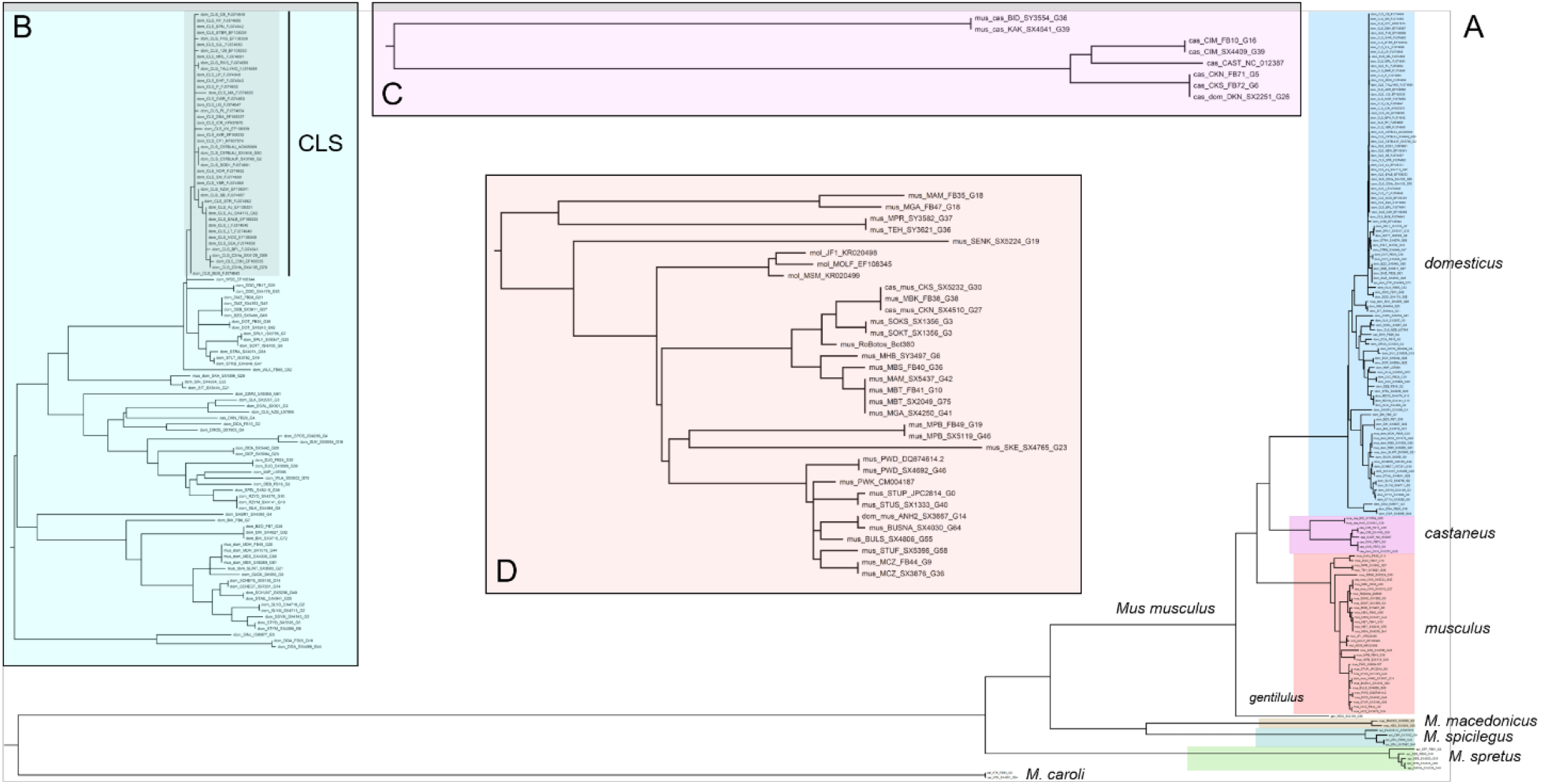
Maximum-likelihood trees for the whole mtDNA dataset (A) and separately for *domesticus* (B), *castaneus* (C), and *musculus* (D) strains. In the *domesticus* tree, the CLS cluster is shaded. The codes for individual samples indicate group, WDS name, mouse ID, and the number of generations of inbreeding. In cases of repeated sequences with different generations, the mouse identification is omitted.

Number of polymorphic sites and nucleotide diversity values can be found in Table 1. We detected 36× more sites and approximately a 100-fold nucleotide diversity in WDS compared to CLS. Interestingly, nucleotide diversity was more than twofold in *musculus*-derived WDS compared to *domesticus*-derived WDS.

**Table 1.** Variation of mitogenomes in the whole dataset.

| Sample analyzed | N | N polymorphic sites | Nucleotide diversity (± SD) |
| --- | --- | --- | --- |
| All strains | 163 | 3907 | 0.03355 ± 0.01594 |
| All CLS | 45 | 108 | 0.00047 ± 0.00025 |
| All WDS | 118 | 3890 | 0.04225 ± 0.02011 |
| *domesticus* WDS | 55 | 915 | 0.00837 ± 0.00404 |
| *musculus* WDS | 36 | 992 | 0.01743 ± 0.00848 |

The numbers of polymorphic sites between individual strains are provided in Supplementary Table S3, while the minimum and maximum numbers of polymorphic sites between the groups are summarized in Table 2. The most significant differentiation was observed between *caroli* and *musculus* mitogenomes, with 2,251 polymorphic sites, accounting for 13.8% of the mitogenome. The non-synanthropic mice (*caroli*, *macedonicus*, *spicilegus*, *spretus*) differed by more than 2,250 polymorphic sites from the synanthropic mice. Differences between house mouse WDS did not exceed 548 polymorphic sites. Despite the *musculus* WDS group having less than half the number of strains compared to the *domesticus* WDS group, variation within the former group was higher as shown in the diagonal of Table 2), confirming the results in Table 1. This difference is primarily attributed to the presence of a 75-bp long insertion in the control region in some *musculus* strains, absent in all *domesticus* WDS. Although some *domesticus* strains possess an 11-bp insert in another part of the control region, which is not found in *musculus*, the 75-bp segment significantly contributes to the higher variation. Indeed, when considering *musculus* WDS without the insert, *domesticus* WDS have up to 40 more polymorphic sites. The same argument applies to the *castaneus* strains.

**Table 2.** Variation in polymorphic sites across the house mouse species/subspecies. Figures in parentheses in the first column represent the numbers of strains. The numbers of polymorphic sites between strains within each group are on the diagonal; maximum counts of polymorphic sites between different groups are listed above the diagonal, and minimum counts of polymorphic sites are provided below the diagonal. The *gentilulus*-derived MGD strain is considered separately here.

| Taxon | *car* | *spr* | *mac* | *spi* | *gen* | *cas* | *mus* | *dom* | CLS |
| --- | --- | --- | --- | --- | --- | --- | --- | --- | --- |
| *caroli* (1) | 0 | 2247 | 2199 | 2187 | 2157 | 2235 | 2251 | 2184 | 2173 |
| *spretus* (4) | 2229 | 98 | 1236 | 1251 | 1209 | 1273 | 1334 | 1220 | 1205 |
| *macedonicus* (2) | 2197 | 1229 | 39 | 835 | 1015 | 1133 | 1168 | 1079 | 1067 |
| *spicilegus* (2) | 2180 | 1243 | 824 | 41 | 1019 | 1126 | 1168 | 1071 | 1057 |
| *gentilulus* (1) | 2157 | 1195 | 1013 | 1016 | 0 | 496 | 545 | 443 | 435 |
| *castaneus* (5) | 2170 | 1193 | 1063 | 1054 | 420 | 269 | 522 | 471 | 465 |
| *musculus* (21) | 2136 | 1183 | 1027 | 1032 | 410 | 375 | 242 | 548 | 534 |
| *domesticus* (48) | 2158 | 1181 | 1047 | 1033 | 421 | 379 | 389 | 149 | 134 |
| CLS (45) | 2164 | 1183 | 1048 | 1046 | 430 | 384 | 393 | 9 | 92 |

A relatively high number of polymorphic sites found within CLS (N=92) was caused by the strain NZB developed at the University of Dunedin in New Zealand ^48^. When the NZB strain was excluded, the level of variation dropped substantially, reaching a maximum of 6 polymorphic sites between C3H and MA strains (Supplementary Table S3). In comparison, the variation within the *domesticus* WDS and *musculus* WDS strains increased by a factor of 1.62 (24.83 when the NZB strain is excluded) and 2.63 (40.33 without NZB), respectively, compared to what was observed in the CLS strain.

The mtDNA dataset included 31 duplicated samples, which were utilized to assess consistency in two aspects: (i) between the newly sequenced and published mtDNAs in CLS and (ii) between mice of the same strain with substantial differences in the numbers of generations since the founding of the strain. While no difference were observed between our and published CLS mitogenomes, inconsistencies between sequences in samples with varying numbers of generations were detected in 12 out of 31 strains (Supplementary Table S4). Remarkably, half of these cases involved introgression between subspecies or even species such as from *spretus* to *domesticus* in STF strain (Supplementary Table S4.A).

#### Prdm9 gene

PRDM9 protein is a histone methyltransferase that initiate meiotic recombination by binding to allele specific DNA sequences via its highly polymorphic zinc finger domain ^49^. To assess its variation, we sequenced the zinc finger (ZnF) domain of the *Prdm9* gene in 98 samples (Supplementary Table S5). This dataset was supplemented with 12 CLS sequences from the study by Kono et al. ^50^. Five *Prdm9* sequences from CLS maintained in Studenec, along with the published sequences from the same CLS ^50^, were used as a sequencing quality control. No differences in duplicated sequences were detected.

By sequencing the *Prdm9* ZnF domain, we identified 48 alleles among the 105 strains analysed (see Supplementary Table S5 for allelic designations, which provide information on the most amino-acid polymorphic sites across all ZnFs, following the nomenclature introduced by Mukaj et al. ^51^). An overall comparison between *M. musculus* WDS and CLS revealed a significant distinction: while only two alleles were found in 12 CLS, as many as 38 alleles were present in 83 *M. musculus* WDS. Allelic diversity detected in the *domesticus* and *musculus* WDS (*A* = 0.51 and *A* = 0.39, respectively) was 3.1–2.4× higher than in the CLS (*A* = 0.17). In contrast to mtDNA, the correspondence between the *Prdm9* gene tree and species tree is weaker. Although most *domesticus*-derived strains, including all analyzed CLS, cluster within a single large group, several of them are interspersed among other clusters. A similar scattered pattern is also observed in WDS representing other (sub)species (Figure 3), with most variable *castaneus* WDS dispersed across the entire phylogenetic tree.

**Figure 3.**
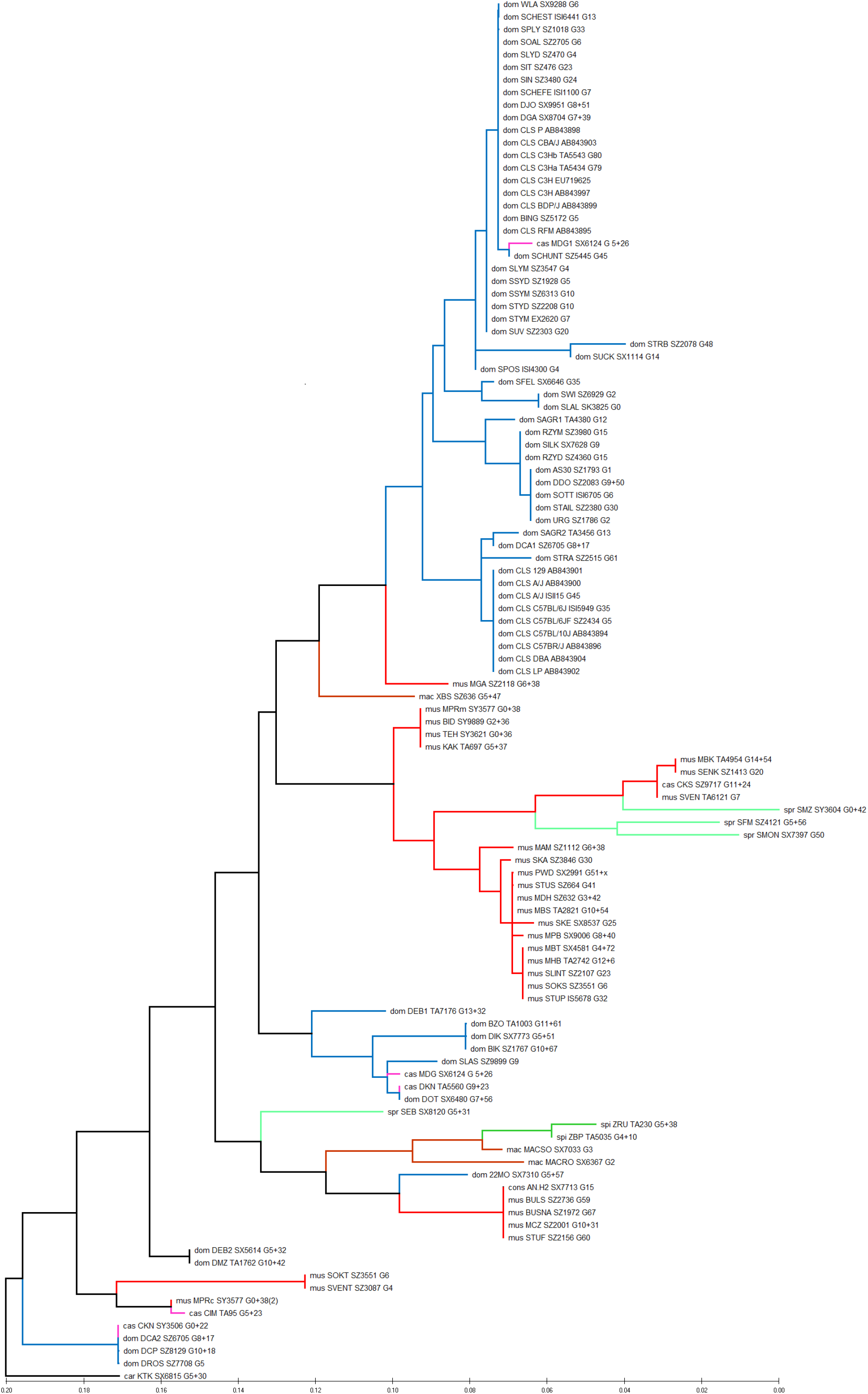
Maximum-likelihood tree of the *Prdm9* gene; blue: *domesticus* WDS (including CLS), red: *musculus*, magenta: *castaneus*, green: *spretus*, brown: *macedonicus*, dark green: *spicilegus*.

#### *Slx*/*Sly* copy number variation (CNV)

We assessed CNV in two highly ampliconic genes, *Slx* (*Sycp3 like X-linked*) located on the X chromosome and *Sly* (*Sycp3 like Y-linked*) on the Y chromosome, in 53 strains (Supplementary Table S6). As shown in Table 3, *musculus* WDS revealed approximately a two times higher CN compared to *domesticus* WDS for both genes. Furthermore, *Sly* CN was, on average, approximately 2.5-fold higher than that of *Slx*. Interestingly, even though both CLS analyzed (C57BL/6J and C3Hb) predominantly possess *domesticus*-like genomes, their *Slx* and *Sly* CNs are considerably higher than those in *domesticus* WDS. The higher *Sly* CN in CLS relative to *domesticus* WDS can be explained by the fact that these strains carry *musculus*-like Y chromosome ^52,53^. However, the average *Sly* CN in CLS is still lower than that in *musculus* WDS (Table 3). Consequently, the two CLS appear intermediate between the *musculus* WDS and *domesticus* WDS clusters when *Sly* CN is plotted against *Slx* CN (Figure 4). It is worth noting that two *musculus* WDS outliers deviate from the overall pattern: MAM and STUS with 21 and 54 *Slx* copies, respectively. These numbers are characteristic of *domesticus* WDS; however, X-linked SNPs in STUS ^9^ and MAM exome ^36^ confirm their *musculus* origin.

**Figure 4.**
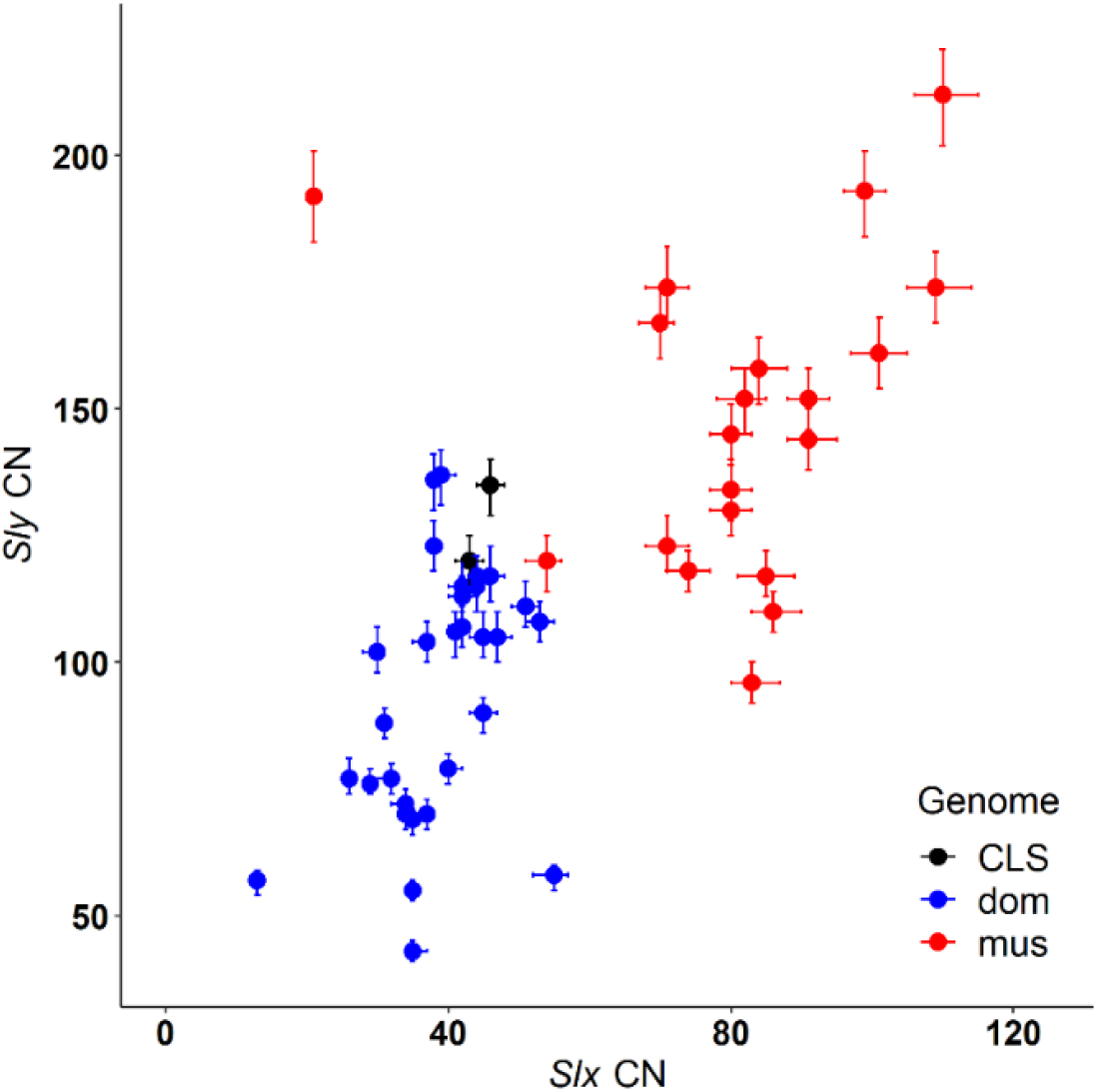
Scatterplot of *Slx* and *Sly* CN in *musculus* WDS, *domesticus* WDS, and CLS. Vertical and horizontal error bars show Poisson distribution-based errors from triplicate measures of *Slx* and *Sly* copy numbers, respectively.

**Table 3.** Mean, minimum and maximum numbers of *Slx* and *Sly* copies in CLS, *domesticus* WDS, and *musculus* WDS; numbers of analyzed strains are in parentheses.

| Ancestry | Mean.*Slx* | Min.*Slx* | Max.*Slx* | Mean.*Sly* | Min.*Sly* | Max.*Sly* |
| --- | --- | --- | --- | --- | --- | --- |
| CLS (2) | 44.5 | 43 | 46 | 127.5 | 120 | 135 |
| *domesticus* (31) | 38.8 | 13 | 55 | 94 | 43 | 137 |
| *musculus* (20) | 81.1 | 21 | 110 | 148.6 | 96 | 212 |

#### Phenotype variability

We assessed phenotypic variation in nine traits recorded in 4,335 mice, representing 84 WDS of *musculus*, *domesticus*, *castaneus*, *spretus*, *spicilegus*, *macedonicus*, *caroli*, and five CLS. Our analysis included only individuals between 65 and 600 days old, since subadult and aged individuals are more likely to be infertile. We measured the following traits: body weight, spleen weight (recorded in 3,080 mice), body length, tail length, weight of both ovaries (measured in 2,051 females), sperm count, the weight of testes, left epididymis, and seminal vesicles (measured in 2,284 males). Sperm numbers were counted in a Bürker hematocytometer, with ten chambers per sample, following the methodology described in Vyskočilová et al. ^54^.

Descriptive statistics of the morphometric data computed separately for males and females revealed high variation both across strains (Supplementary Table S7) and the groups (Supplementary Table S8). ANOVA detected significant differences between the groups for all measured traits, including sex*group and sex*strain interactions for external measurements (Supplementary Table S9). It is worth noting that males of Palearctic non-synanthropic species (*macedonicus*, *spicilegus* and *spretus*) displayed substantially higher testis weight and sperm count values, while ovary size showed lower differentiation among females across all groups (Supplementary Figure S1).

The analysis of morphological data, restricted to CLS, *domesticus* WDS, and *musculus* WDS revealed substantial differentiation between the groups both for males and females. Out of 39 comparative tests for the nine traits, only five were insignificant (4× for CLS and *domesticus* [males: testicular and epididymal weight; females: tail length and ovary weight], 1× for CLS and *musculus* [males: sperm count]) (Supplementary Table S10). Individual-based analyses of the morphospace, defined by the projection of two variables, confirmed a higher divergence of the three groups in external traits than in the reproductive characteristics (Figure 5A-C, and 5a-c). In general, the contribution of CLS to the overall morphological variation was negligible, ranging from 0.0% (testis weight vs. sperm count) to 0.1% (body weight vs. relative tail length) (Figure 5a and 5c). In contrast, males and females of WDS exclusively contributed 57.7% and 53.7%, respectively, to the total variation defined by body weight and relative tail length; the remaining proportion of the morphospace was shared among CLS and WDS. This pattern differs from the reproductive traits, where most of the variation is shared between all three groups in males (59.9% in testis weight vs. sperm count, Figure 5C and 5c) and between *domesticus* and *musculus* and all three mouse groups in females (each contributing 46.8% and 44.2%, respectively, in the morphospace defined by body weight and ovary weight [data not shown]).

**Figure 5.**
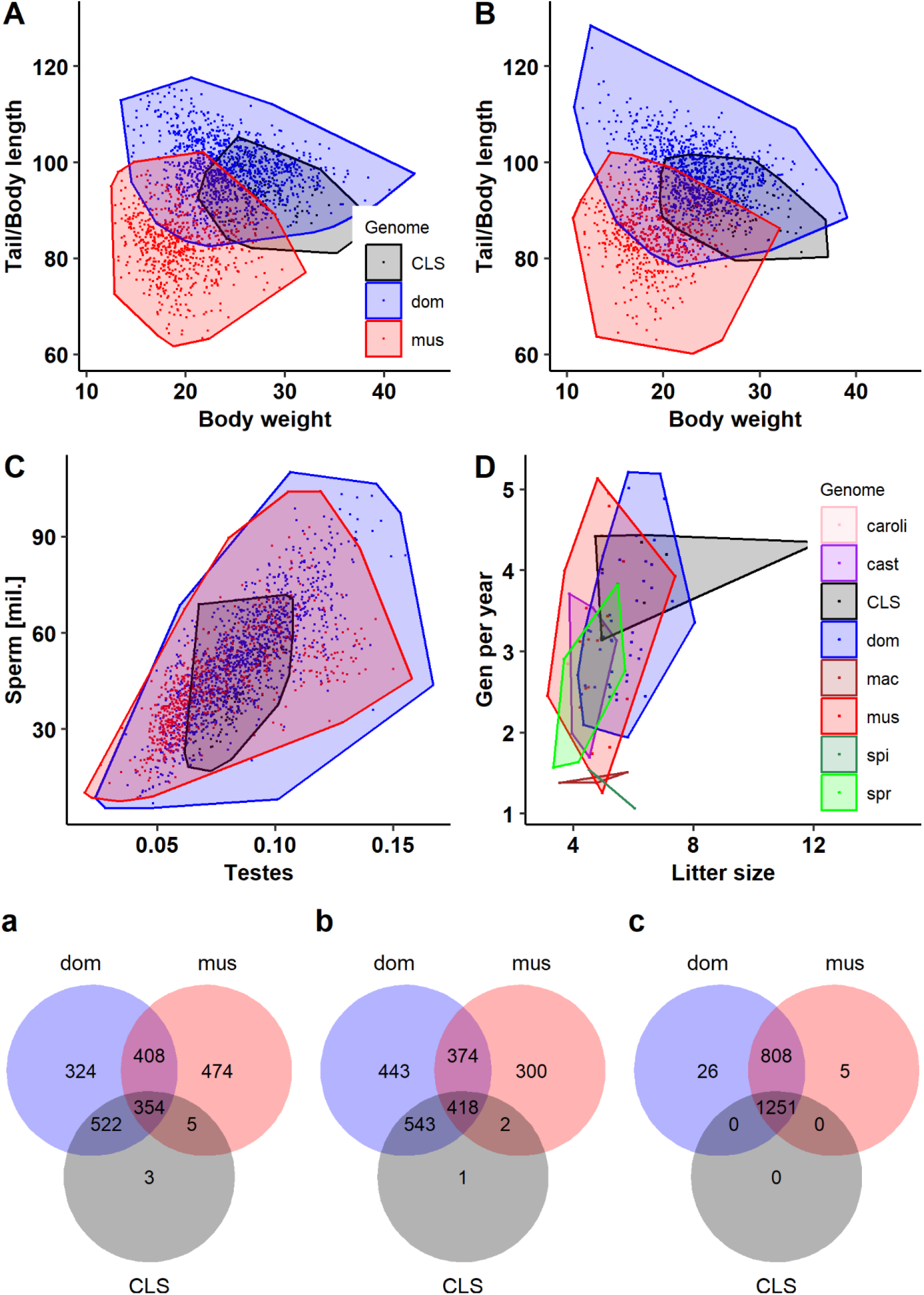
Morphospaces defined by biplots of body weight vs. relative tail length (100*body length/tail length) in males (A) and females (B); testis weight vs. sperm count (C); and mean litter sizes vs. number of generations delivered per year (D). Small lettered Venn diagrams provide the numbers of overlapping and non-overlapping individuals in *domesticus* WDS (dom), *musculus* WDS (mus), and CLS for corresponding graphs labelled with capital letters (A–C).

#### Reproduction performance

Reproductive ability was estimated in 87 WDS and 8 CLS. In total, the dataset consisted of 90,077 offspring born to 8,298 mothers in 17,049 litters recorded in Studenec studbooks between 2000 and 2023. Reproductive performance was characterized by litter size, newborn mortality (calculated as the proportion of stillborn or cannibalized mice across all litters), and the number of generations produced per year. We also estimated the time since a WDS was established from wild progenitors until completing 20 generations of strict brother-sister mating, i.e., the generation at which a strain of mice can be considered inbred ^55^. These data were summarized across the groups and individual strains.

Table 4 lists reproductive parameters summarized for each group of mice (note that, based on exome data, *gentilulus* is grouped with the *castaneus* MDG strain ^36^). Excluding *caroli* and *spicilegus* WDS with fewer than three strains per taxon, litter size, mortality, and the number of generations delivered within a single year showed significant differences among the groups (ANOVA, P < 0.005 for all variables). CLS were the only stocks that could produce, on average, more than six pups per litter and four generations per year (Table 4). Furthermore, along with *domesticus* WDS, CLS were characterized by newborn mortality below 10%. In this regard, CLS exhibit reproductive characteristics that are highly suitable for maintaining and efficiently breeding, reaffirming their significance as an ideal model in biomedical research. The non-synanthropic species exhibited the lowest reproductive performance among all the groups. This fact can have significant implications when planning and preparing experiments using newly derived strains: while achieving the desired level of inbreeding would take less than five years in CLS or their derivates (consomics, congenes), and approximately seven years in *musculus*-derived WDS. However, the process could extend to about 14 and 16 years for *macedonicus* and *spicilegus* strains, respectively.

**Table 4.**
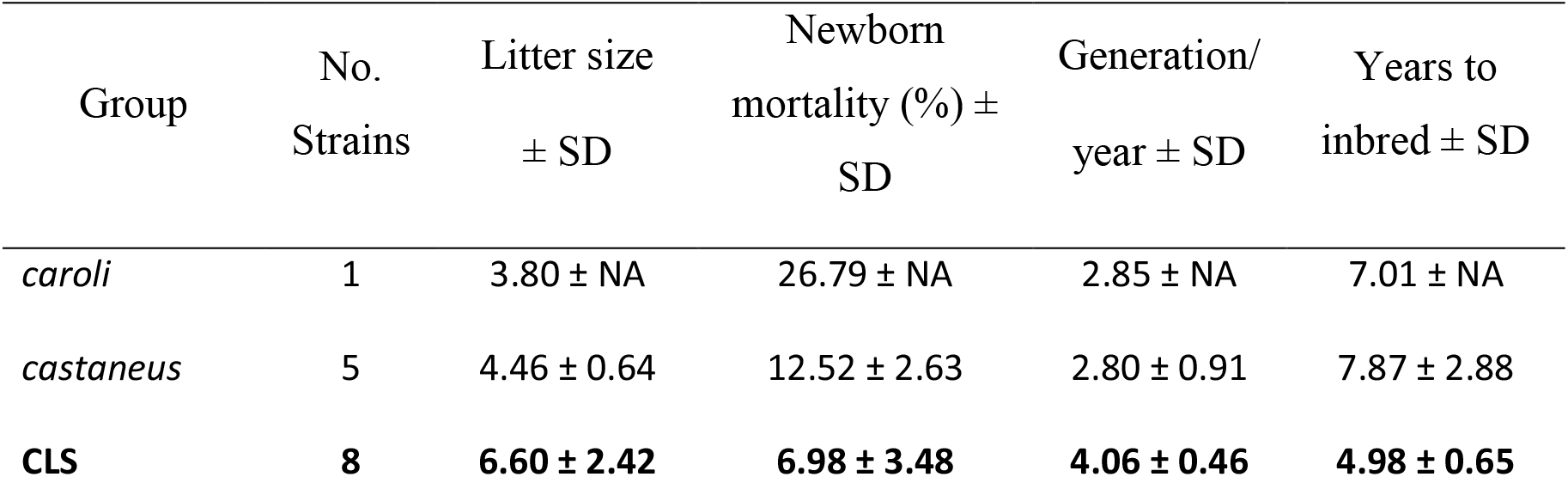
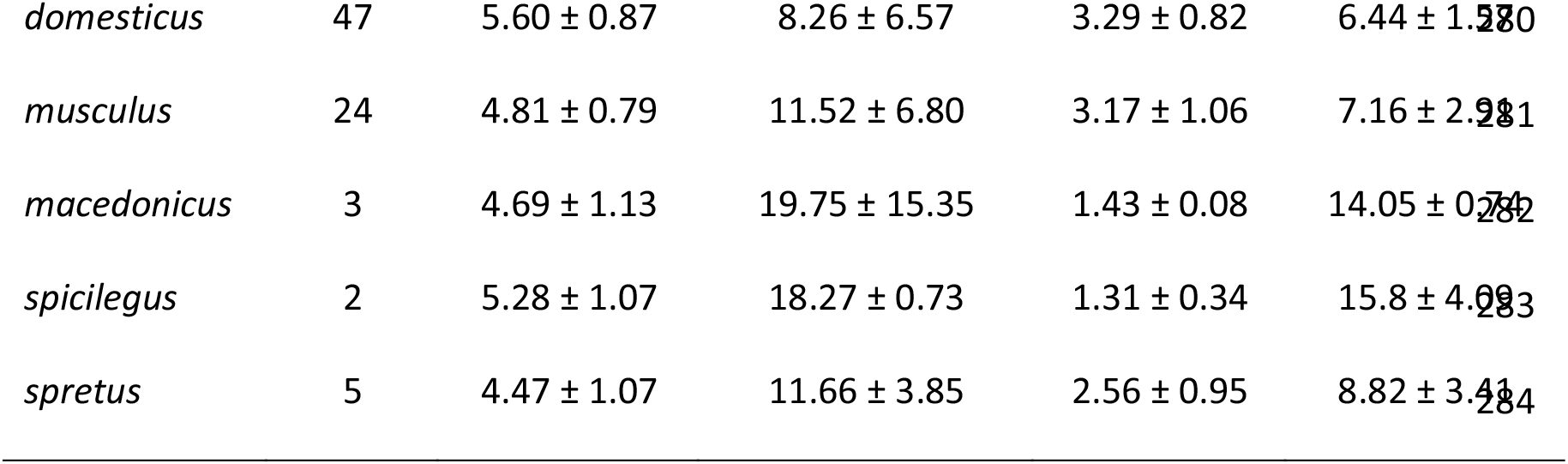
Reproductive characteristics of WDS and CLS. ‘Years to inbred’ represents the average number of years required to achieve mouse strains in the 20^th^ generation of strict inbreeding. Bold figures indicate the highest scores.

High variation in the reproductive parameters is also present among individual strains (Supplementary Table S11). For example, the difference between the minimum and maximum litter sizes is 3.92-fold, ranging from 3.11 (*musculus*: MDH) to 12.20 (CLS: CD-1). CD-1 (outbred mice derived from a group of Swiss albinos) was the only strain with an average litter size exceeding 10 (as indicated by the rightmost outlier in Figure 5D). Similarly, higher inter-strain variability was observed in the rate of reproduction, with a 4.87-fold difference between the minimum and maximum numbers of generation produced per year, ranging from 1.07 (*spicilegus*: ZBP) to 5.21 (*domesticus*: BING). Alhough it cannot be tested, these traits may have both genetic and environmental components, and, as a result, their values can vary between different laboratory settings.

Two of the three reproductive parameters differed between *musculus* WDS, *domesticus* WDS, and CLS (ANOVA: litter size: P=0.0008; newborn mortality P=0.0539, generations/year: P=0.0441). Pairwise comparisons between these three groups showed significant differences in three of the nine comparisons (Supplementary Table S12): *musculus* WDS had a lower litter size than *domesticus* WDS and CLS. Additionally, CLS were the only strains capable of producing over four generations per year; however, they exhibited significant differences only from *musculus* WDS (Table 4, Supplementary Table S12). Despite our best efforts to maintain all strains in optimal conditions, some strains stopped reproduction in various generations. The extinction rate observed across 99 strains was 23.2% (Supplementary Table S1).

## Discussion

Any model organism would benefit from incorporating natural variation, to assess how accurately laboratory stocks represent the species they are meant to emulate and provide insight into molecular processes and their variability. Such an integration would significantly enhance the relevance of model systems for the study of complex trait variations, including diseases ^56^. In this study, we examined genetic and phenotypic variation across nearly 100 wild-derived mouse strains to comprehend the complexity of the vertebrate model. The data were compared to CLS, which are often taken as representatives of the house mouse in comparative studies e.g., ^57,58,59^.

Genetic and phenotype variation was estimated using mitochondrial and nuclear DNA sequences, copy number variation of two ampliconic sex-linked genes, and 11 morphological and reproductive traits. As expected, genetic variation was significantly higher in WDS compared to CLS, and its magnitude increased with divergence time. Similarly, the phenotypic diversity preserved in WDS substantialy exceeded and differed from that observed in classical laboratory mice. Both datasets, i.e. genetic and phenotypic, are congruent, suggesting that CLS harbour only a tiny fraction of the house mouse variability and hence do not adequately represent mice in natural populations. Incorporating variation from WDS can, therefore, enhance the biological reality of the model. In conclusion, we advocate for including the variation preserved in WDS, which holds great potential to enhance the effectiveness of the house mouse model not only in evolutionary studies but also in biomedical research.

### WDS versus CLS

Given their different sources and histories, higher genetic variability in WDS compared to CLS is expected. However, the magnitude of this difference, as summarized in Supplementary Table S13, is stunning. For example, WDS mitogenomes harbour approximately 90 times higher nucleotide diversity than CLS. None of the mitogenomes is shared between CLS and WDS. The allelic diversity in the *Prdm9* gene, which plays an important role in *musculus*/*domesticus* hybrid male sterility ^35,60^, is approximately three times higher in *domesticus* and *musculus* WDS than in CLS. Of the two *Prdm9* alleles present in CLS, only one (dom3) was detected among WDS, while the other allele (dom2) is private for CLS (but present in wild *domesticus* in North America and North Europe – Emil Parvanov and Jiří Forejt, personal communication 2021, ^61^). *Prdm9* primarily defines the positions of recombination hostpots during meiosis^62^. Consequently, the presence of only two alleles in CLS generally limits the mapping of quantitative trait loci in laboratory crosses. The presence of 35 *Prdm9* alleles in *musculus* and *domesticus* WDS, along with the substantial inter-strain/inter-subspecific variation documented here, may challenge this limitation and provide rich material for testing genotype-phenotype associations.

Copy number variation is an important driver of genome and phenotype evolution ^63^. It is worth noting that the proportion of the autosomes and the X chromosome affected by CNV was found to be lower in CLS (estimated size of 1.7 Mb, representing 0.065 % of the mouse genome) compared to WDS (3.8 Mb, representing 0.14 % of the genome) ^64^. In this study, we analyzed CNV in *Slx* and *Sly* genes, whose disrupted balance has been associated with infertility and sex ratio bias in house mice ^65–68^. Recently, it has been shown that an arms race between these genes significantly affects the dynamics of the hybrid zone between *musculus* and *domesticus* ^27^. Interestingly, the number of gene copies of CLS *Slx* and *Sly* appeared intermediate between the *musculus*-derived and *domesticus*-derived strains (Figure 4; Table 3). Since both CLS analyzed in this study possess predominantly *domesticus*-like autosomal genes and *musculus*-like Y chromosomes ^9,52,53^, we should expect X-linked *Slx* CN and Y-linked *Sly* CN to fall within *domesticus* and *musculus* range, respectively. In the context of the WDS/CLS comparison, it is important to note that because of their hybrid origin, CLS resemble *domesticus* populations close to the hybrid zone center, possessing introgressed *musculus* Y chromosomes ^27,28,69^ rather than genetically pure *domesticus* populations. Moreover, due to the presence of chromosomal fusions and whole-arm reciprocal translocations, the western house mouse (*domesticus*) has become a well-known model for the study of karyotype variation ^70,71^. While only the standard karyotype with 2*N* = 40 acrocentric chromosomes occurs in CLS, 12 of 40 *domesticus* WDS karyotypes display reduced diploid numbers ranging between 22 and 38 chromosomes (Supplementary Table S1).

Morphological variation mirrors genetic variability. It is noteworthy that CLS are almost entirely embedded within morphospace defined by WDS. On the other hand, when comparing CLS with *domesticus* and *musculus* WDS across individual traits, we observe significant differences in most external morphological and reproductive traits (Supplementary Table S13). As anticipated, due to their genomic composition, the morphological differentiation between CLS and *musculus* WDS is higher than that between CLS and *domesticus* WDS (Supplementary Table S13). Possibly because of long-term artificial selection in captivity, CLS outperform WDS in terms of reproductive parameters (Table 4), which highlights their suitability as models for biomedical research. The phenotypic differentiation between WDS and CLS observed here in congruent with the study of Takada et al. ^72^ comparing 10 WDS derived from *domesticus*, *castaneus*, and *molossinus* subspecies and one CLS (C57BL/6). In that study, significant differences were found in 62—68% of 21 measured morphological and physiological traits ^72^.

In summary, the genetic data corroborate previous analyses that have found higher diversity in wild mice compared with CLS e.g. ^9,36,53,72,73^. Simultaneously, *Slx* and *Sly* CN, along with the numerous significant differences in external morphology and reproductive characteristics, suggest CLS are not simply domesticated representatives of *M. m. domesticus*, despite the overwhelming portion of *domesticus* genome they carry ^9^. It seems the genetic and morphological differences documented here reflect not only the complex history of CLS but also their long-term adaptation to laboratory conditions, influenced by genetically conditioned behavioural changes fixed during domestication ^25^. In recognizing their unnatural genetic constitution, CLS have been suggested to be called *‘Mus laboratorius’* ^31^ or *‘Mus gemischus’* ^74^. Here we added a phenotypic dimension to this perspective.

### WDS versus wild mice

While we revealed significantly higher variation in WDS compared to CLS, an important question to consider is to what extent WDS truly represent wild mice For instance, in comparing CNV as a quantitative trait, we found that differentiation in *Slx* and *Sly* copy number between *domesticus* and *musculus* WDS mirror what has recently been reported from the European house mouse hybrid zone ^27^. When examining mtDNA, the most comprehensive data has been gathered from analyzing *cytochrome b* (*mt-Cytb*) and D-loop sequences. In *domesticus*, seven (*mt-Cytb*) and 11 (D-loop) haplogroups have been described ^75–77^, five *mt-Cytb* haplogroups were found in *musculus* ^78^. WDS displayed six haplogroups in *domesticus* and the same number in *musculus* strains. These numbers suggest that while WDS preserve variation from the major representatives of deeply diverged haplogroups they may struggle to capture natural variation on local scales within these haplogroups. For example, an analysis of whole mitogenome sequences in 98 wild mice, mostly from the eastern Palearctic, revealed the presence of 90 haplotypes ^47^. Similarly, while we identified 48 variants in the *Prdm9* gene in 99 WDS, 207 different alleles were identified in 453 wild and wild-derived animals ^50,61,79,80^. These figures highlight that while WDS capture variation from various lineages, they may lack the discriminatory power to differentiate individual variation within those lineages.

The documented diversity preserved in WDS can be a valuable resource for investigating genotype-phenotype variation in evolutionary contexts. For example, the observed variation of *Prdm9* alleles in WDS can provide insight into our understanding of the recombination machinery associated with the dynamics of formation and erosion of *Prdm9*-dependent binding sites ^81^. Furthermore, negative interactions of specific alleles in intersubspecific mouse hybrids can result in meiotic arrest and male sterility ^35,51,60,82,83^. As mentioned earlier, the WDS described in Supplementary Table S1 preserve approximately 25% of currently known natural *Prdm9* allelic variation. With this limited number of alleles, we can still observe the extent and intensity of negative interactions among 36 alleles in 1296 F1 hybrids within and between *domesticus* and *musculus* WDS. Importantly, since some alleles are shared among multiple strains (e.g., the sterility-causing allele dom3 in 8 *domesticus* WDS), it becomes possible to assess the effects of genetic background and the evolution of sterility in natural populations.

While there are numerous genetic comparisons between wild-derived mouse strains (WDS) and wild mice, comparative morphometric studies between these two groups are scarce. For example, a study employed four sperm size parameters to assess whether WDS can serve as a proxy for exploring evolutionary processes related to post-copulatory selection ^84^. For this purpose, 28 wild-caught and four WDS derived from *domesticus* and *musculus* subspecies were utilized. The subspecies exhibited significant differences in sperm head length and midpiece length, and these differences were consistent for wild mice and wild-derived strains when pooled over genomes. However, when the inbred strains were individually analyzed, their strain-specific values sometimes significantly deviated from subspecies-specific values obtained from wild mice. This study, therefore, suggests that future experiments should involve a larger number of strains to account for natural variation and avoid confounding results due to reduced variability and founder effects within individual stocks ^84^. In this respect, we believe that the documented genetic variation in the resources of WDS presented in this paper (and maintained in other laboratories) can effectively capture the majority of genetic and morphological diversity observed in their wild counterparts.

### WDS cross-contamination

An important consideration when using wild-derived mouse strains (WDS) for modelling evolutionary processes is ensuring that these strains represent natural, not artificially introduced, genetic variation. In a study by Yang et al. ^9^, traces of intersubspecific contamination were reported in the WDS under investigation. They proposed that introgression in these WDS could have resulted from a combination of cross-contamination from CLS, gene flow in the wild (in WDS captured near hybrid zones), or breeding with other wild-derived mice in laboratory settings. We analyzed potential cross-contamination in our WDS panel by comparing mitogenomes of the same strains sampled at different times during their existence. Inconsistencies in these duplicated samples were observed in 38.7% of 31 cases (Supplementary Table S4), indicating the presence of introgression from other genomes. It is worth noting that some cross-contaminations can be attributed to the breeding scheme used to maintain mouse stocks. For instance, certain WDS were initially kept through breeding among distantly related pairs (outbreeding) at the University of Montpellier, a practice that inherently maintains much genetic polymorphism. Subsequently, these outbred stocks turned to brother-sister mating and are now considered inbred. However, without complete information on the original parental individuals, it remains unclear which alleles were fixed in subsequent generations. This change in the breeding scheme can explain at least the intrasubspecific shifts between mtDNA haplogroups. On the other hand, intersubspecific mtDNA shifts, like the presence of a *domesticus* mitogenome with an 11-bp insertion in the D-loop of two *musculus* WDS from Bulgaria, may suggest unintended cross-contamination. The fact that this +11bp variant is widespread in northern Germany and Scandinavia and matches a haplotype of a Danish *domesticus* WDS further supports its artificial origin in the Bulgarian strains. Similarly, two *castaneus* strains from Kenya (CKN and CKS), were found to share the same *musculus* haplotype as in the *musculus*-derived MBK strain from Bulgaria. A possible explanation is that during the inbreeding process, which inevitably leads to the fixation of deleterious variants, there is a strong advantage for the offspring of an unintentional interstrain cross restoring fertility. Such events can potentially occur in mouse facilities where stocks are kept in the same room.

To conclude this section, although we have detected traces of artificial introgression in 15.2% of the analyzed 79 WDS (Supplementary Table S4), the majority of these strains remain a valuable source of natural variation. They have the potential to complement and expand research that has traditionally relied heavily on CLS. Given the time-consuming nature of strict inbreeding, which can span from 6.5 to 17 years to obtain an inbred strain, we believe it is essential to initiate the documentation and characterization of mouse resources developed and maintained in various laboratories worldwide. Making these resources available to the scientific community can have a significant impact. In a prospective view, since WDS can provide reproducible genotypes similar to CLS, they can be integrated into Phenome projects once they have been transferred to specific-pathogen-free facilities.

### Relevance of WDS to human and biomedical research

From an evolutionary perspective, the genotype-phenotype variation preserved in mouse WDS is undoubtedly very important. However, it can also serve as a surrogate for studying human populations in both evolutionary and biomedical contexts. The synanthropic bond between house mice and humans began to evolve approximately 10,000 years ago with the onset of human agriculture ^85,86^. This association has allowed mice to inhabit new human-associated ecological niches and spread across the globe ^31,32,87^. The colonization of new environments has driven adaptations to varying local conditions, as observed along a latitudinal gradient on the eastern coast of North America ^20^. These adaptations to local conditions are expected to result in global intrasubspecific and intersubspecific divergence. Indeed, an analysis of more than 150 whole-genome sequences of wild mouse populations sampled worldwide revealed that a significant fraction of genetic variation is private to individual populations ^88^, which can limit GWAS studies of their associations. Interestingly, the analysis also detected strong signals of positive selection in many genes associated with human diseases ^88^.

Among the traits examined in this study, mtDNA holds significant importance due to its known role in causing several inherited diseases in humans ^89^. CLS are frequently utilized in these studies ^90^. Here, we document that the variability in mtDNA within our mouse WDS exceeds that observed in humans. To illustrate this point, we compared our finding to a dataset comprising 560 maternally unrelated human individuals of European, African, and Asian descent published by Elson et al. ^91^. A pairwise haplotype comparison within this human dataset revealed a maximum difference of 260 polymorphic sites, which is similar to the variation observed within intrasubspecific *musculus* and *domesticus* WDS, but lower than what was found in intersubspecific *domesticus*/*musculus* WDS comparisons (548 polymorphic sites; see Table 2). Such comparisons hold the potential to become a cornerstone of biomedical research, where the haplotype diversity detected in WDS can be harnessed, for example, in the context of tissue transplantation. The introduction of new nuclear DNA–mtDNA interacting systems may accelerate the onset of metabolic disorders in the recipient organism as reviewed in ^92^. Understanding the mechanisms governing competitive mtDNA segregation and identifying which tissues can tolerate heteroplasmy is essential for making more accurate predictions in these interactions. Using a mouse model, it was demonstrated that mtDNA segregation in heteroplasmic mice depends on the genetic divergence between a donor and a recipient and is also tissue-dependent, implying potential complications in human therapies^93^. Therefore, the concept of haplotype matching has been proposed as an approach to mitigate these issues in the context of mitochondrial replacement therapies ^93,94^.

A similar pattern emerges when comparing the variation in the *Prdm9* gene, which also governs the distribution of recombination hotspots in humans ^95^. Alleva et al. ^96^ studied *Prdm9* allelic diversity in 720 individuals from seven worldwide human populations and detected 69 alleles (i.e., approximately 0.10 per individual), compared to 38 alleles observed here in 83 *M. musculus* WDS (i.e., approximately 0.46 per strain). While our focus here has been on individual representatives of mouse genomes, recent whole-genome analyses of laboratory stocks (including three WDS) have identified that the most variable regions of the mouse genome are enriched with genes relevant to disease and infection response ^97^. Returning to humans, a new pangenome human reference map that aligns 47 genome assemblies from genetically diverse individuals promises to enhance our understanding of genomics and our ability to predict, diagnose and treat diseases ^16^.

### Relevance of WDS for preclinical testing

Drug discovery and development is a long, costly, and high-risk process spanning over 10–15 years, with an average cost of over $1–2 billion for each new drug to gain approval for clinical use ^98^. Despite rigorous optimization of drug candidates during the preclinical stage, nine of ten candidates allowed to proceed to clinical studies fail during one of three phases of clinical trials in the drug approval process ^99^. New approaches, such as machine learning or open-source cross-sector collaboration, are being employed to reduce the failure rate of medications in preclinical or clinical settings. However, significant effort is still directed toward enhancing the rigor and reproducibility of testing ^100,101^.

Due to high demands on reproducibility, CLS are the dominant animal models used in drug testing and the development of therapies for human disease. On the other hand, the fact that mouse WDS exhibit dramatically higher genetic and phenotypic variation than CLS and that their genetic variation is comparable to that found in human populations can greatly enhance their relevance in medical research, for example, in preclinical testing of the effectivity of biomolecules in *in vivo* pharmacology. When WDS are integrated into preclinical biomolecule testing and complement the battery of tests performed on CLS, the effectiveness of molecules can be assessed across a broad spectrum of genotypes. Molecules that exhibit specific responses within a limited range of genetic variability can be identified and excluded from further evaluation. Consequently, the risk of failure in clinical trials can be significantly reduced, potentially saving up to 90% of the associated costs.

## Material and Methods

### Mice

All mice are housed in a conventional breeding facility (i.e., without nanofilter barrier or specific-pathogen free condition) of the Institute of Vertebrate Biology, Czech Academy of Sciences, in Studenec. They are maintained under standard conditions, with a light/dark regime of 14/10 hours, with temperatures of 23 ± 1 °C during summer (April-September) and 22 ± 1 °C during winter (October-March), respectively. The relative humidity is maintained within 40-70%. Mice have access to food pellets (Myška 1, VKS Podhledští Dvořáci, Hamry, Czechia) and tap water *ad libitum*. Mice are weaned at 20 days of age and housed in pairs in EURO IIL cages with a floor area of 530 cm^2^ made of transparent polycarbonate (Tecniplast, Varese, Italy). The cages are equipped with bedding material (sifted sawdust, Happy Horses, Martinsberg, Austria), shredded paper for nest building and nest houses made of red transparent polycarbonate (Tecniplast). The facility is authorized for the use of experimental animals (licences 61974/2017-MZE-17214 and MZE-50144/2022-13143), as well as for the breeding and supply of experimental animals to third parties (62065/2017-MZE-17214 and MZE-50151/2022-13143). These licences are in compliance with the corresponding regulations and standards of the European Union, as specified in Council Directive 86/609/EEC.

In June 2023, we acquired additional seven WDS consisting of 1 *musculus* (Kazakhstan), 4 *domesticus* (France, Germany, two from Iran), 1 *castaneus* (Taiwan) and 1 *spicilegus* (Slovakia). These stocks were previously maintained at the Max Planck Institute for Evolutionary Biology in Plön, Germany. While six of them were initially kept in outbred mating scheme in Plön, they have been subjected to strict brother-sister mating since their transfer to Studenec. More detailed information about these stocks can be found in Harr et al. ^19^. However, due to recent commencement of their strict inbreeding, these stocks could not be included in the analyses.

### mtDNA

Genomic DNA was isolated from frozen (−80 °C) or alcohol-preserved muscle or spleen tissues using DNeasy Blood & Tissue Kits (Qiagen). High-quality DNA aliquots were next-generation sequenced at the Edinburgh Genomics facility using HiSeq X technology. Two-paired 150 bp reads were merged and mapped against the C57BL/6 mitogenome (GeneBank NC_005089) ^12^ using the Geneious Prime software (Biomatters: www.geneious.com). The fine-tuning iteration was set to 5. In the cases where the presence of the 75-bp insertion in the control region ^102^ was indicated by a peak of increased read coverage in the respective region, the sample was re-mapped against *de novo* assembled mitogenome of an *M. m. musculus* individual (Bot360) from Botosani, Romania, which is known to carry the insert ^103^. The average coverage was 225 ± 151 reads per sample (individual data are available in Supplementary Table S2). Consensus sequences were generated with the sequence matching threshold set to 65%, and ‘N’ was called in a sequence when the coverage was less than 3. These data were supplemented with 49 published WDS and CLS mitogenomes ^12–14,104–106^.

All sequences were aligned using Clustal Omega ^107^ implemented in the Geneious Prime (Biomatters Ltd., www.geneious.com). Maximum likelihood phylogenetic trees were inferred using the GTR+G+I model ^108^ selected with jModelTest ^109,110^. Six discrete categories ^111^ were used to approximate the continuous gamma distribution. In addition, an extensive subtree pruning and regrafting procedure ^112^ was applied to improve searching for the best tree. The weakest stringency of optimization with respect to branch lengths and improvements in log-likelihood values (branch swap filter) was used to maximize the explored search space. The MEGA11 software ^113^ was employed for the analysis.

### *Prdm9* sequencing

We followed the protocols published by Buard et al. ^61^ and Kono et al. ^50^. The ZnF array was PCR amplified using PrimeSTAR HS DNA Polymerase (Takara Bio) and primers Prdm9-F (TGAGATCTGAGGAAAGTAAGAG) and Pdrm9-R (TCCTGTAATTGTTGAGATGTGG); 20 μl of the total volume included 30 ng of genomic DNA and 0.5 μM of each primer. The PCR conditions were as follows: after 30 s at 98 °C, 28 cycles were carried out, including 10 s at 98 °C, 15 s at 63 °C, and 2 min at 68 °C. The PCR product’s size was determined using electrophoresis in 1% agarose gel (Seakem). In cases where mice were heterozygous and carried alleles of different lengths, the corresponding bands were excized from the gel and purified using ethanol precipitation. A second round of PCR was performed using sequencing primers: Prdm9seqF (CTCAGAACAGGCCAGACAACA) and Prdm9seqR (TTGTTGAGATGTGGTTTTATTGCT). Sanger sequencing was conducted by Eurofins Genomics (Olomouc, Czechia) from both ends of the purified PCR products, using the sequencing primers. It is important to note that many PCR products could not be sequenced up to their ends in both directions due to the repetitive nature of ZnF arrays. In such cases, sequencing was repeated until high-quality electropherograms were obtained.

Assembly of the forward and reverse sequences and the translation of DNA sequences into amino acids were performed using Geneious Prime. To define individual *Prdm9* alleles, triplets of amino acids located in the most variable positions, specifically positions −1, +3, and +6 of each ZnF, were utilized, as described by Oliver et al. ^114^. The sequences were aligned with ClustalW. Maximum likelihood tree was inferred in MEGA11 ^113^ using default values.

### *Slx/Sly* copy number variation

The digital droplet PCR (ddPCR) method ^115^ was employed to estimate copy number in two ampliconic genes, *Slx* (*Sycp3 like X-linked*) and *Sly* (*Sycp3 like Y-linked*), in 53 strains. A custom-designed PrimeTime qPCR assays (IDT^®^, Coralville, Iowa, USA) were used. Two assays were run in duplex reactions, where each well contained one primer pair and probe designed for the target gene (either *Slx* or *Sly*) and one for the *Tert* gene. *Tert* was used as a reference since it is known to be consistently present in two copies in diploid organisms ^116^. The assay sequences and PCR conditions were identical to those used in Baird et al. ^27^.

The ddPCR reactions were carried out in the Genomics Core Facility at the Central European Institute of Technology (CEITEC) in Brno using the QX100/200 Droplet Digital PCR System (Bio-Rad, Hercules, CA, USA). For each sample, measurements were performed in triplicate. Quantasoft^TM^ Software (Bio-Rad) was used to estimate the error and merge the three obtained values into a single outcome representing the number of gene copies.

### Statistical analyses

The analysis of variance (ANOVA) was used to detect variability in the morphological traits. The factors compared in the models were group, strain, sex, and their interactions. The analyses were based on two models. The first model tested global differentiation within the whole dataset comprising eight groups, 90 strains, and, where appropriate, sex. The second model was confined to the three most representative groups: *musculus* WDS, *domesticus* WDS, and CLS (3,968 mice: 2,096 males and 1,877 females). This dataset was lower for splenic weights and consisted of 1,459 males and 1,359 females. All statistical analyses were conducted in the R statistical language R Core ^117^ and run in the R Studio environment RStudio ^118^. Pairwise comparisons between group means were tested using the emmeans package (https://CRAN.R-project.org/package=emmeans).

Two reproductive traits deviated from normal distribution. Litter size was normalized by excluding one outlier with an exceptionally large litter size (CD-1 with 12.2 young in a litter). For the mortality rate, a normal distribution was achieved using a square root transformation. The third variable describing the rate of reproduction (the average number of generations delivered per year) displayed normal distribution.

Moreover, we aimed to determine the proportion of variation in a morphospace defined by length or mass variables specific to each group and the proportion shared between groups. For each group, we first computed the subset of points lying on the convex hull of the set of points specified in the biplot projection of two length/mass variables using the ‘hull’ algorithm ^119^. We then calculated the numbers of points falling into individual polygons and their intersections using the ‘point.in.polygon’ function from the ‘sp’ package in R. These analyses included three groups of mice: *musculus* WDS, *domesticus* WDS, and CLS. The ‘ggvenn’ R library was used to visualize data in the form of Venn diagrams. Regarding morphometric traits, we used data for body weight and tail/body length ratio, which is known to be higher in *domesticus* than in *musculus* ^120,121^. We also separately analysed morphological variation defined by testis weight and sperm count in males and ovary weight and body mass in females. Reproductive ability was investigated in a similar way for all eight groups.

## Supporting information

Supplementary Material Table S1. List of strains kept in mouse repository in Studenec.

Supplementary Material Table S2. List of strains analysed for whole mitochondrial genome.

Supplementary Material Table S3. Pairwise SNPs differences between strain mitogenomes.

Supplementary Material Table S4. Mitogenomes: intersubspecific and intrasubspecific clade shifts in duplicated samples.

Supplementary Material Table S5. List of strains analysed for the Prdm9 gene.

Supplementary Material Table S6. Copy number variation at Slx and Sly genes in strains.

Supplementary Material Table S7. Descriptive statistics of morphological traits in strains in males and females.

Supplementary Material Table S8. Descriptive statistics of morphological strains in mouse groups for males and females.

Supplementary Material Table S9. ANOVA tests of morphological traits in all strains.

Supplementary Material Table S10. Pairwise contrasts between CLS, domesticus and musculus strains for phenotype traits.

Supplementary Material Table S11. Reproduction parameters of WDS.

Supplementary Material Table S12. Statistical testing for contrasts in reproductive parameters between CLS, domesticus and musculus strains.

Supplementary Material Table S13. Overall summary of pair-wise differences between CLS, domesticus and musculus WDS.

## Acknowledgements

We would like to express our gratitude to Stuart Baird for providing hybrid indices of strains. The construction of the breeding facility in Studenec was made possible through funding from the Czech Academy of Sciences. The long-term housing of mouse strains was supported by the Institute of Vertebrate Biology, with additional funding from Czech Science Foundation Grants No. 16-23773S, 17-04364S, 19-19056S, 21-28491S, 22-29928S as well as CAS within the program of the Strategy AV 21. We appreciate the access to the National Grid Infrastructure MetaCentrum, provided under the programme ‘Projects of Large Infrastructure for Research, Development, and Innovations’ (LM2010005). Core Facility Genomics of CEITEC Masaryk University (supported by the NCMG research infrastructure LM2023067 funded by MEYS CR) is gratefully acknowledged for providing the equipment used for CNV estimations. We would also like to thank Christine Pfeifle and Diethard Tautz for generously donating the strains derived at the Max-Planck-Institute.

## Author Contributions

J.P. designed the study, collected mice, was responsible for morphological and reproductive performance analyses, and wrote the initial version of the manuscript. Ľ.Ď. collected mice, derived chromosomal and morphological data. Z.H. assessed CNV. J.K. designed statistical analyses, T.A., A.B. D.Č., J.GdB, and K.J. sequenced mtDNA. H.H., JanaP., I.M. and I.P. took care of mice. L.R. heads breeding facility, took care of mice and genotyped mice for the presence of the *t*-haplotype. A.O. and F.B. were responsible for a mouse repository in Montpellier, B.V.B. collected mice and genotyped Y chromosome. J.F. is responsible for a mouse repository in Prague and wrote the manuscript. M.M. analysed sequence data and wrote the manuscript. P.K. sequenced the *Prdm9* gene, designed routine genotyping of strains at diagnostic markers and wrote the manuscript. All authors have read and approved the final version of the manuscript.

## Data records

Sequences of 114 complete mtDNA genomes were submitted to GenBank with accession codes GB000X–GB00XX.

Znf array sequences of the *Prdm9* gene of 98 mice can be accessed at GenBank under OQ054372– OQ054469.

WDS are registered at Mouse Genome Informatics http://www.informatics.jax.org/strain/summary. Further data on this mouse repository can be obtained at https://housemice.cz/en/strains/.

## Supplementary data

**Supplementary Material FigureS1.**
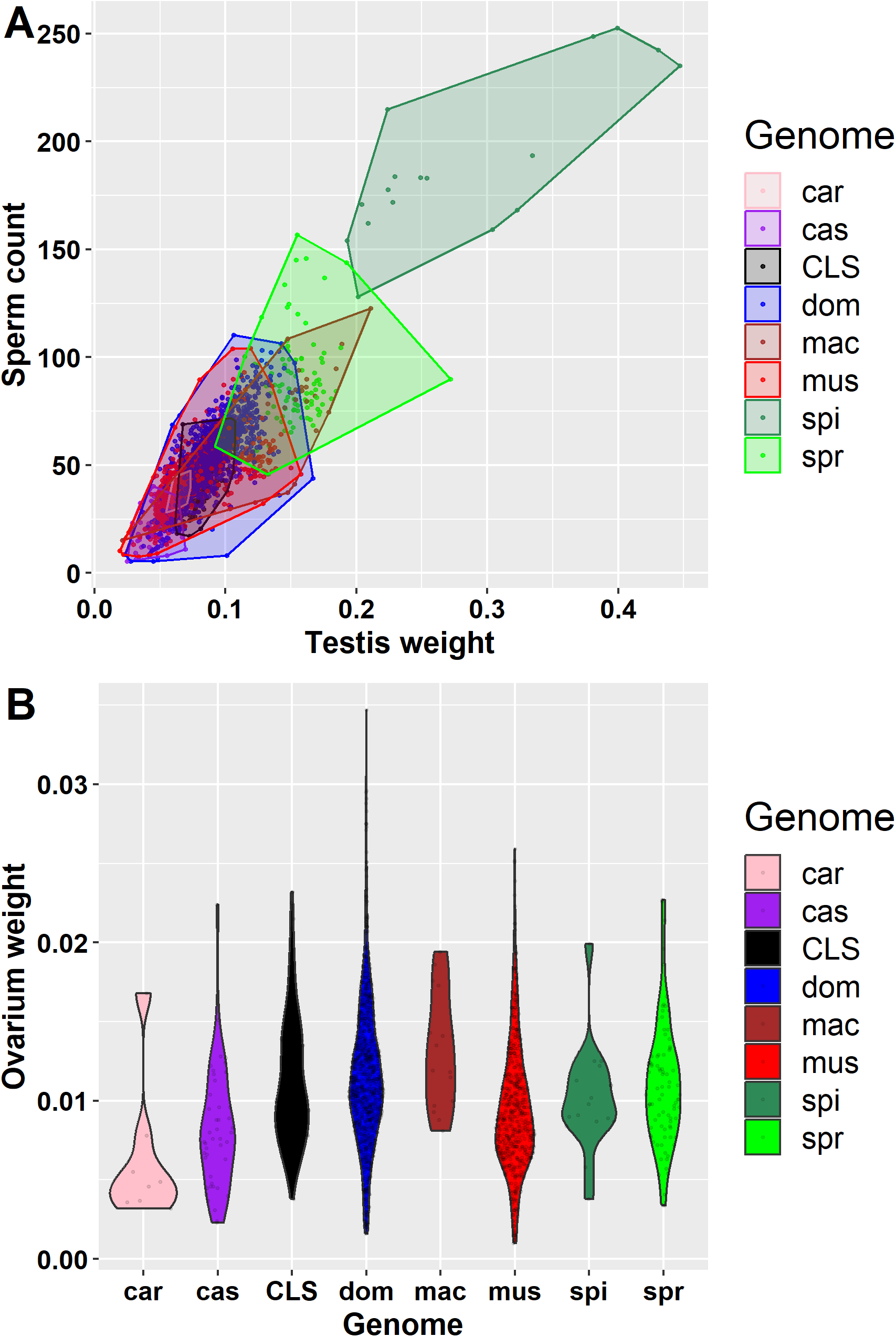
Variation in reproductive traits of males (A) is higher than in females (B). Data span in CLS does not exceed morphospace defined by variation in WDS.

